# A tree-planting decision support tool for urban heat island mitigation

**DOI:** 10.1101/821785

**Authors:** Zoey Werbin, Leila Heidari, Sarabeth Buckley, Paige Brochu, Lindsey Butler, Catherine Connolly, Lucila Houttuijn Bloemendaal, Tempest D. McCabe, Tara Miller, Lucy R. Hutyra

**Author notes:** These authors contributed equally to this work (Zoey Werbin, Leila Heidari, Sarabeth Buckley). These authors also contributed equally to this work (Paige Brochu, Lindsey Butler, Catherine Connolly, Lucila Houttuijn Bloemendaal, Tempest D. McCabe, Tara Miller, Lucy R. Hutyra).

## Abstract

Heat poses an urgent threat to public health in cities, as the urban heat island (UHI) effect can amplify exposures, contributing to high heat-related mortality and morbidity. Urban trees have the potential to mitigate by providing substantial cooling, as well as co-benefits such as reductions in energy consumption. The City of Boston has attempted to expand its urban canopy, yet maintenance costs and high tree mortality have hindered successful canopy expansion. Here, we present an interactive web application called “Right Place, Right Tree - Boston” that aims to support informed decision-making for planting new trees. To highlight priority regions for canopy expansion, we developed a Boston-specific Heat Vulnerability Index (HVI) and present this alongside maps of summer temperatures. We also provide information about tree pests and diseases, suitability of species for various conditions, land ownership, maintenance tips, and alternatives to tree planting.

## Introduction

In a changing climate, urban areas are facing hotter, longer summers with more extreme heat events and increased heat-related mortality and morbidity [1,2]. In Boston, for example, the heat-related mortality rate “may more than triple to 10.5 per 100,000 people under a moderate emissions reduction scenario or reach as high as 19.3 per 100,000 under the business-as-usual emissions scenario” by the 2080s [3]. Cities are often significantly warmer than surrounding rural areas, acting as “heat islands” due to heat-absorbing construction materials and low tree canopy cover [4,5]. Urban tree canopy substantially reduces summer temperatures and air-conditioning costs, yet tree canopy is often unequally distributed along class and race line [6,7]. This makes canopy expansion an important tool for building climate resilience during hot periods, as well as improving social equity in cities [5,7].

The City of Boston has markedly lower canopy cover than other New England cities, with trees present on about 23% of land [8]. Boston leadership has attempted to expand its urban canopy, without much success [9]. In 2007 Boston set a now-abandoned goal to plant 100,000 trees, called “Grow Boston Greener,” yet urban canopy may actually have decreased since the goal was set [10,11]. Although Boston officials are aware of tree-planting as a powerful climate mitigation and environmental health practice, high tree mortality and deferred maintenance hinder efforts to expand Boston’s canopy [3,12,13].[11–13]. The city of Boston – like many other cities without a current master plans for tree planting – would benefit from new tools to aid in urban canopy planning.

On another dimension, regional Heat Vulnerability Index-based recommendations have not been implemented into Boston’s planting strategies. An HVI helps identify regions more susceptible to heat-related morbidity and mortality based on the demographic profiles of people who live there, for example older adults or younger children who are more susceptible to heat[14]. A 2015 Tufts University study suggested priority regions for tree planting in Boston based on a Heat Vulnerability Index (HVI) [14]. Another component of an HVI is the land use in different regions, which can exacerbate or improve urban heat islands. A 2015 Tufts University study suggested priority regions for tree planting in Boston based on a Heat Vulnerability Index (HVI) which considers social vulnerability, adaptive capacity, and physical variables [14]. HVI has been shown to have accurately describe heat-related mortality, and can be used to develop effective, targeted interventions [15].

The City of Boston and Boston University (BU) partnered in 2018 to address questions posed by municipal decision-makers around tree planting. An interdisciplinary team of PhD students at Boston University working at the nexus of environmental sciences and public health was challenged with how to maximize the benefits of future tree-planting efforts, while taking into consideration areas and populations most vulnerable to the health effects of heat, as well as ensuring tree survival and feasibility. We took advantage of the recent advances in the availability of public data that can inform heat vulnerability, as well as developments in technology for creating interactive web tools to distribute statistical analyses, to create a decision support tool called *Right Place, Right Tree* | *Boston*. In this paper, we describe the analyses and recommendations that fed into a new decision support tool, as well as the tool itself. We also describe our engagement with stakeholders to improve the tool’s usability and practicality, as well as tips for reproducing this tool for other cities.

## Materials and Methods

### Overview

Through this collaborative effort, the team produced a decision support tool to encourage successful planting: Right Place, Right Tree | Boston. This tool implements the following decision-making framework: Step 1) Choose a region for planting trees: identify priority census tracts based on heat or HVI, as well as planting feasibility; Step 2) Learn about regional considerations: consider region-specific factors such as asthma and pest prevalence, existing canopy, land ownership, and community partners; Step 3) Choose the right tree species: reduce tree mortality and maintenance costs by choosing a tree that suits the site conditions well; and Step 4) Keep your tree healthy: explore resources for tree maintenance, education, and legal matters. In Steps 1 and 2, HVI assessment, planting feasibility analyses, and land temperature are reported at the census tract level for consistency with available health data and previous Boston HVI work [20]; our census tract analyses are therefore complemented by a species selector (Step 3) that incorporates site-specific considerations.

### Heat Vulnerability Index

The Boston HVI was calculated based on methods used by Nayak et al. [14], briefly summarized as follows: the 13 variables identified by Nayak et al. [14] to be related to heat vulnerability were extracted from the American Communities Survey (ACS) 2009-2013 5-year estimate [16] and the 2011 National Land Cover Database (NLCD) [17]. Variables included sociodemographics at a census tract level (i.e. percentage of the population that is Hispanic, black, foreign born, speak English less than “very well,” with income below poverty level, over 65 years of age, over 65 years of age and living alone, age 18–64 years with a disability, age 18– 64 years and unemployed). Also included were percentage houses built before 1980, density of housing units per square mile, percentage land with high building intensity areas and of open undeveloped areas. We used SAS 9.4 [18] to perform rotated Principal Components Analysis (PCA) to assign weighting factors to each variable, retaining components that met the eigenvalue and interpretability criteria. Broadly the three components retained were related to the following dimensions: 1) sociodemographic, 2) urbanicity, and 3) older age and isolation. [14]. We then multiplied each variable by PCA weight and standardized the weighted variables to have a mean of 0 and a standard deviation of 1, and further categorized them (scores 1-6) based on standard deviation for a neighborhood from the observed mean, with the least vulnerable category (less than 2 standard deviations below the mean) assigned a score of 1, and the most vulnerable category (more than 2 standard deviations above the mean) assigned a score of 6. These factor scores were then summed to an overall HVI score, where a high score indicated high heat vulnerability across one or more dimensions.

### Planting Feasibility

Our planting feasibility metrics rely on a dataset produced by the University of Vermont Spatial Analysis Lab (UVM SAL). The UVM SAL created a geospatial land cover dataset using 2013-2016 imagery of Boston [19]. This dataset included mutually-exclusive classifications such as buildings, impervious surfaces, vegetation (grass/shrub), and tree canopy. Our analysis focused on the opportunities for immediate tree planting, impervious surfaces were not considered to be “potential” canopy. Potential canopy was defined using the vegetation land cover class [19]. Data were reported in square footage at the block group spatial scale, which was aggregated to the census tract level. Existing and potential canopy is reported as both total square footage and percentage of total area; both metrics are informative due to variation in census tract size. Processing of geospatial data was done in ArcGIS Pro, ArcMap 10.6, and SAS 9.4. Long-term mean summertime land surface land temperature [20] and our HVI metric were used to prioritize census tracts for intervention. Canopy expansion was the recommended intervention for census tracts with high planting feasibility; alternative solutions were recommended for census tracts with low feasibility.

### Species Selector

Once census tracts with high planting feasibility *and* high temperature or heat vulnerability were identified, we built a dataset of tree species, based on the 34 tree species approved for urban planting by the City of Boston. We compiled tree characteristic data that could be used to filter tree species, including canopy spread, light requirement, resistance to breakage, site type (i.e. street or park/yard), and pollen allergenicity levels. Trees with high potential to reduce land surface temperatures are highlighted with an icon. Data on canopy spread, light requirements, and resistance to breakage were extracted from species fact sheets produced by the Forest Service [21]. Species recommendations for streets or parks/yards were produced by the Arnold Arboretum [22]. Allergens were included [23] because they can trigger asthma symptoms, creating an environmental health concern; planting high-allergen trees can therefore counteract some of the equity improvements that trees otherwise represent. Tree heat-reduction potential was derived using estimates of species transpiration rates and species leaf area, and was produced by i-Tree, a peer-reviewed software suite developed by the U.S. Department of Agriculture (USDA) Forest Service [24]. For the trees recommended based on the above conditions, we report additional information from the Forest Service fact sheets that can inform decision making, such as tree height, growth rate, pest and disease vulnerability, soil condition requirements, drought tolerance, and pruning requirements.

### Right Tree, Right Place Boston Decision Tool Development

The decision-making tool was developed using the Shiny web framework for R 3.5.1 [25,26]. App framework was created using the *shinydashboard* R package [27]. Maps were created using census data downloaded through the *tigris* R package [28] and visualized using the *leaflet* R package [29]. The species selector was created using simple filtering from the curated tree database using the *DT* R package [30]. Code and data required to generate the application are hosted at https://github.com/zoey-rw/Boston_trees (DOI: 10.5281/zenodo.3515227).

## Results and Discussion

### Recommendations for heat reduction through urban canopy expansion

Trees bring a number of significant benefits to a region, including heat-reduction, pollution mitigation, and stormwater runoff absorption [20,31,32]. Because tree planting initiatives often underserve low-income communities and communities of color [33,34], canopy expansion efforts should aim to reduce inequities by planting trees with social and environmental health considerations in mind. These considerations are captured by our Heat Vulnerability Index (HVI). With a goal of reducing the urban heat island effect, however, planting in the hottest areas may also be a practical approach. We identified priority areas for canopy expansion at the census tract level, based on either high surface land temperature (“heat”) or high HVI. There was little overlap between high heat and high HVI tracts, so the two priority “scenarios” are reported separately (SI Table 1). For each scenario, we report high planting feasibility tracts for which canopy expansion is recommended (Fig 1).

**Fig 1.**
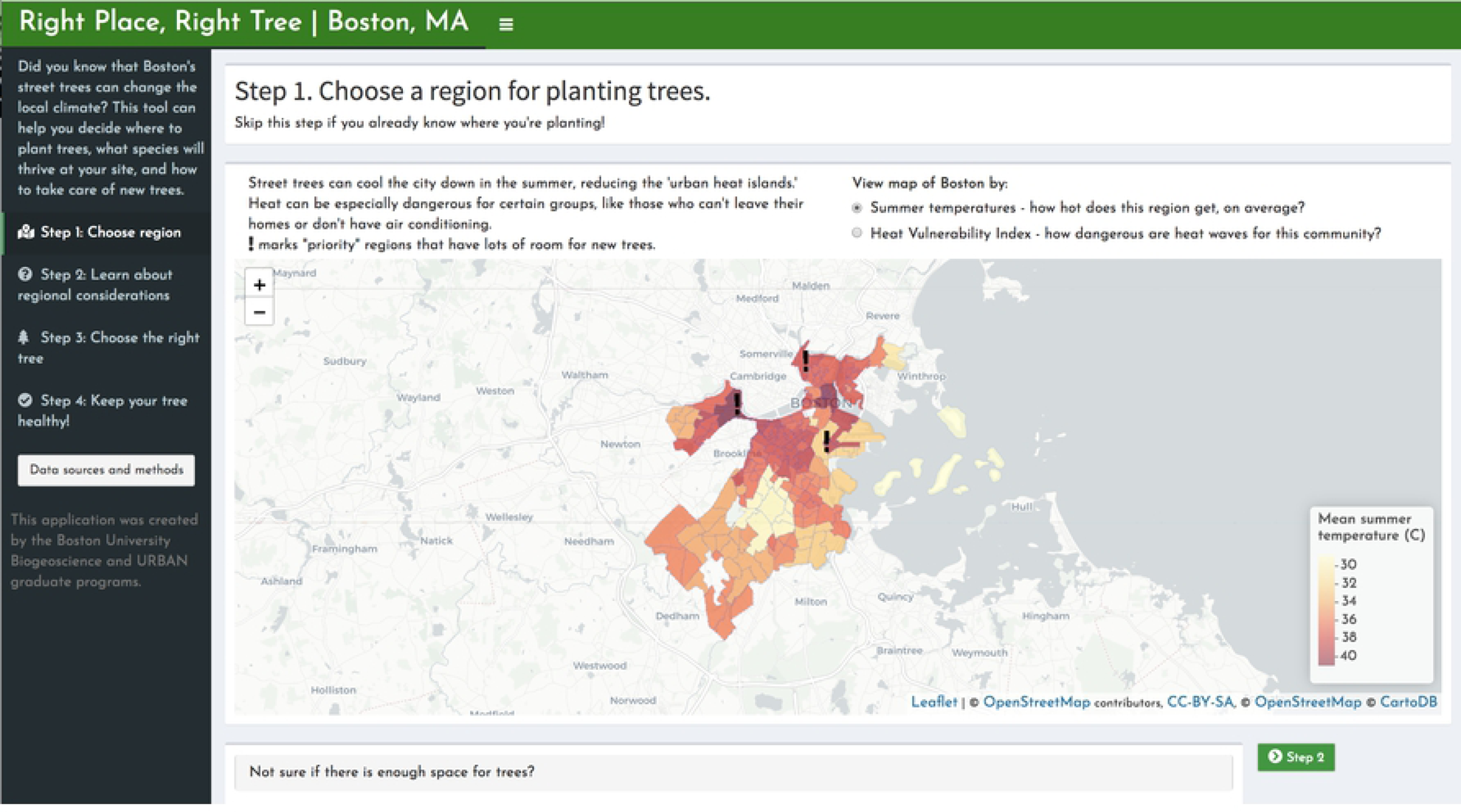
“Step 1: Choose a region for planting trees.” The first page of the interactive decision-support tool recommends identifying a region (census tract) within Boston for planting trees. This page displays maps of summer temperatures as well as Heat Vulnerability Index (HVI), and clicking on the map displays the neighborhood name. Below these maps, an expandable section addresses the potential for canopy expansion for each census tract, and provides a list of resources for alternatives to tree-planting that may also counter the urban heat island effect.

Alternative solutions for heat mitigation include approaches that reduce heat in individual buildings, such as opening cooling centers and retrofitting existing buildings with cool roofing materials. It is also possible to remove impervious surfaces, which would open up more space for potential vegetation. The city could follow New York City’s lead in beginning a depavement campaign on public lands [35], and there are local grassroots efforts to depave various private properties [36], which could be subsidized or encouraged by the city. Green roofs and green walls are additional ways that plants can be integrated with the existing cityscape and buildings. Green roof gardens are particularly known for their ability to cool urban spaces [37], and can help address stormwater issues by capturing rainwater and removing it through evapotranspiration [38]. Green spaces like green roofs and walls have community benefits: both increase the aesthetic appeal of urban spaces, and green roofs in particular can provide new spaces for community events, opportunities for engaging with the process of growing plants, and even food if a rooftop farm is established [39]. Incorporating public-benefit features such as these could be required of all new buildings or buildings undergoing major renovations. This has successfully been incorporated into the policies of cities including Toronto and Chicago [40].

Other recommendations for urban heat reduction include a review of maintenance approaches, approved trees, and public-private partnerships. The current approach includes professional maintenance guarantees for 2 years, after which is when tree mortality is highest [13]. Expanding the duration of planting maintenance contracts, or enlisting more community support for watering, may reduce the deaths of young trees. A high priority should also be placed on protecting older, larger trees, which contribute a disproportionate amount of environmental benefits [13].

Boston’s list of 34 approved trees also includes species with potential invasive qualities (e.g. *Acer campestre*), high VOC emissions (e.g. *Nyssa sylvatica, Liquidambar styraciflua*), and low benefits for biodiversity (e.g. *Ginkgo biloba*); we therefore recommend that this approved tree species list be carefully reviewed and updated. Opportunities exist for public-private partnerships for tree planting and maintenance, such as Cambridge’s “Back of Sidewalk” program, in which the city plants trees on private land up to 20 ft from public sidewalks; this allows trees to be planted in conditions where they are more likely to thrive [41]. Tree-planting practices that involve community engagement may also be more likely to succeed [34].

### Decision-support Tool

Priority area analyses and other relevant information have been compiled in a web application called *Right Place, Right Tree* | *Boston*. The app is organized into four tabs for the following decision-making framework: 1) Identify priority areas based on heat or HVI (Fig 1). On a map of Boston, census tracts with high canopy expansion feasibility and high heat or high HVI values are highlighted. Below this map, users can view the potential for canopy expansion in each census tract. 2) Once an area of interest is chosen, area-specific factors can be evaluated (Fig 2). A clickable map of Boston returns higher-resolution data on asthma prevalence, community partners, and land ownership. 3) Choose a tree species that fits the site conditions well (Fig 3). A species selector allows users to filter trees by site-related preferences, such as light availability, available area, site type, allergenicity, and maintenance requirements. 4) Keep your tree healthy, using a collection of resources for city officials and residents, including city-produced guides, pest sighting updates, and links for requesting maintenance via Boston’s 311 service (Fig 4).

**Fig 2.**
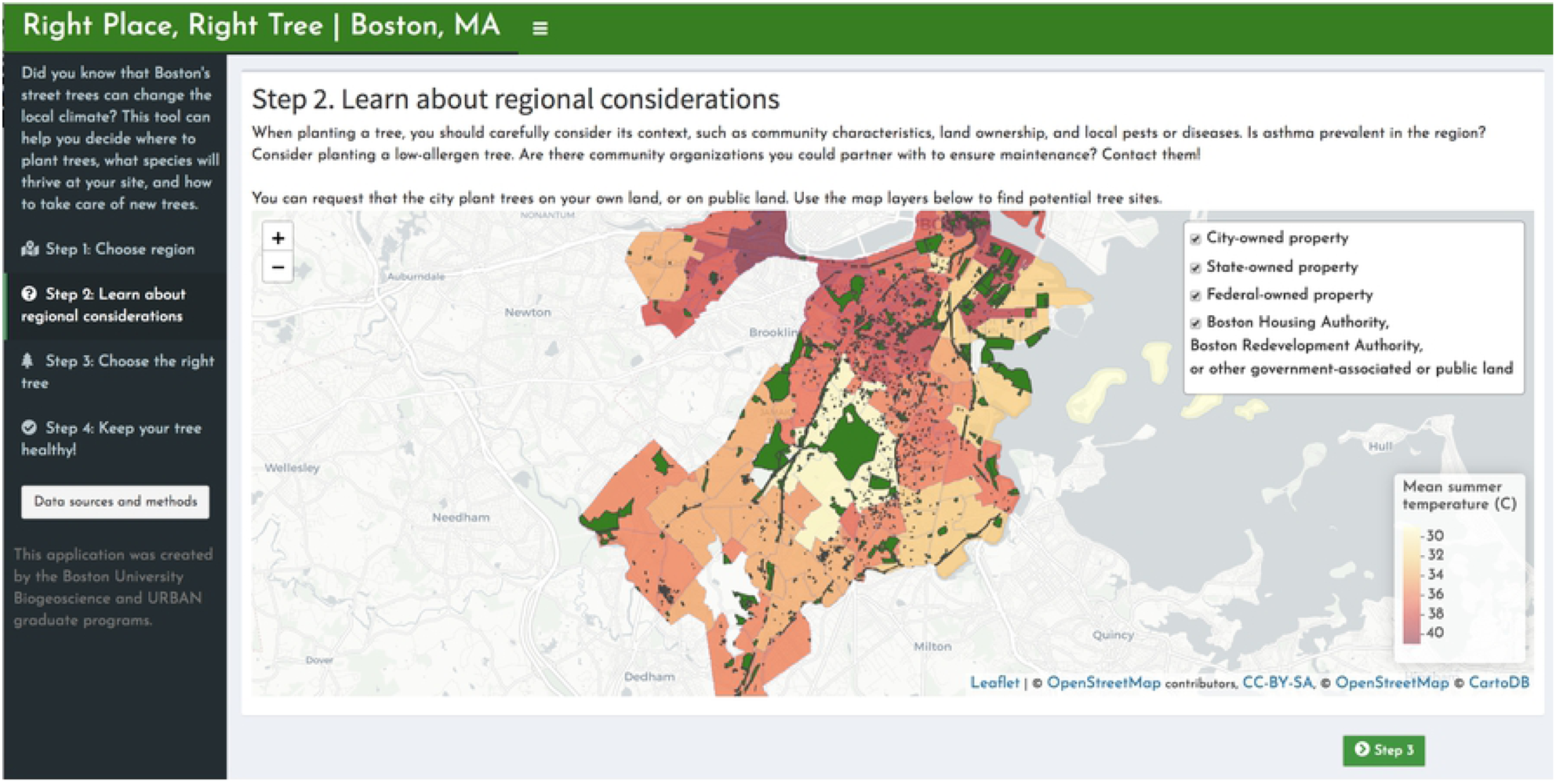
“Step 2: Learn about regional considerations.” The second page of the tool displays a map of Boston with additional information that may influence decision-making, such as pest sightings, property ownership, and names of community organizations that may assist with planting or maintenance.

**Fig 3.**
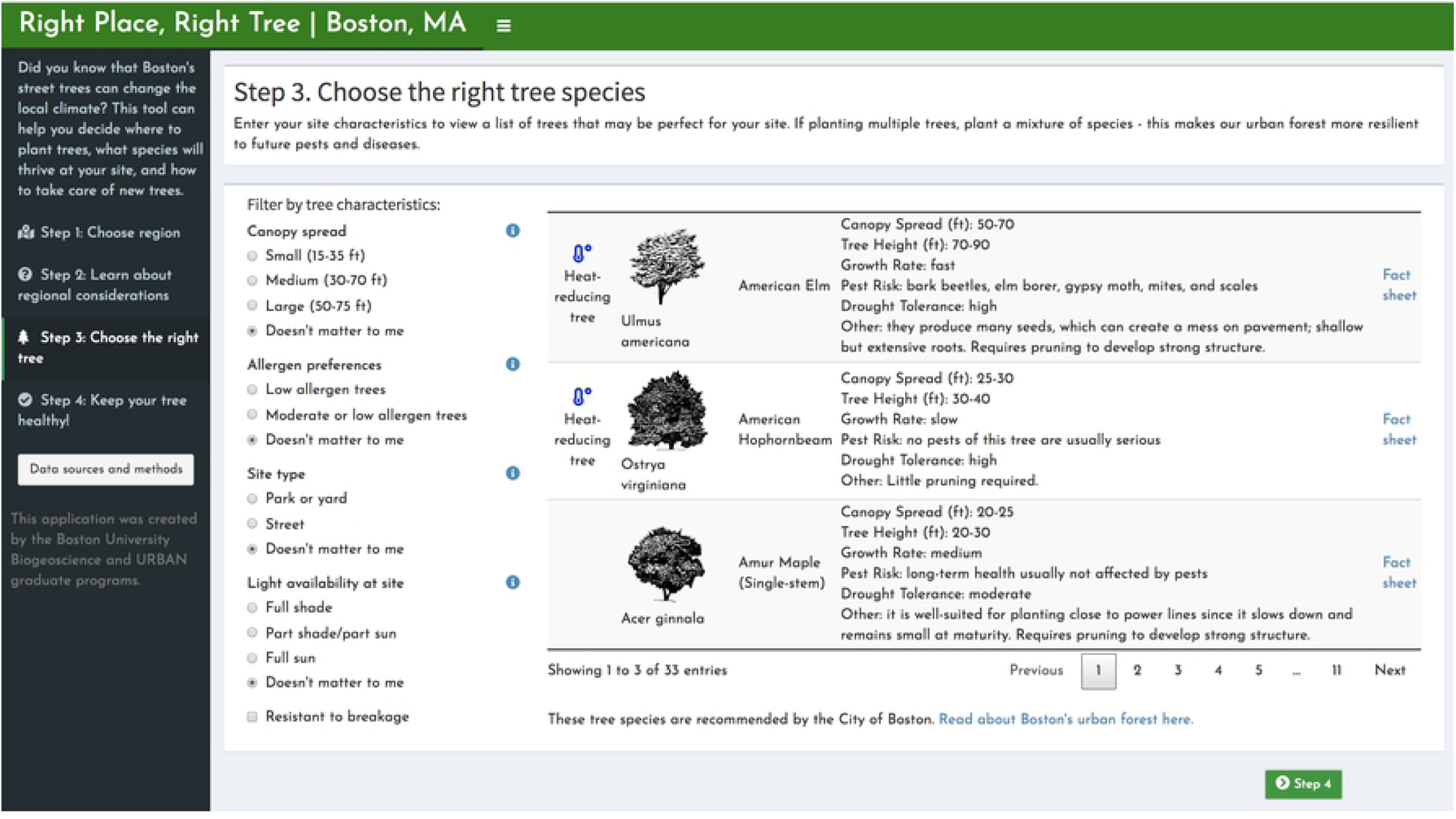
“Step 3: Choose the right tree species”. The third page of the tool allows for the user to use preferences and site characteristics to filter a list of tree species approved by the City of Boston. Choices can be selected on the left and the trees that meet all specifications appear to the right. Each listing includes a small image, basic size and growth information, a link to a full fact sheet, and an icon that indicates if a tree has particularly high potential for heat-reduction (based on transpiration rates).

**Fig 4.**
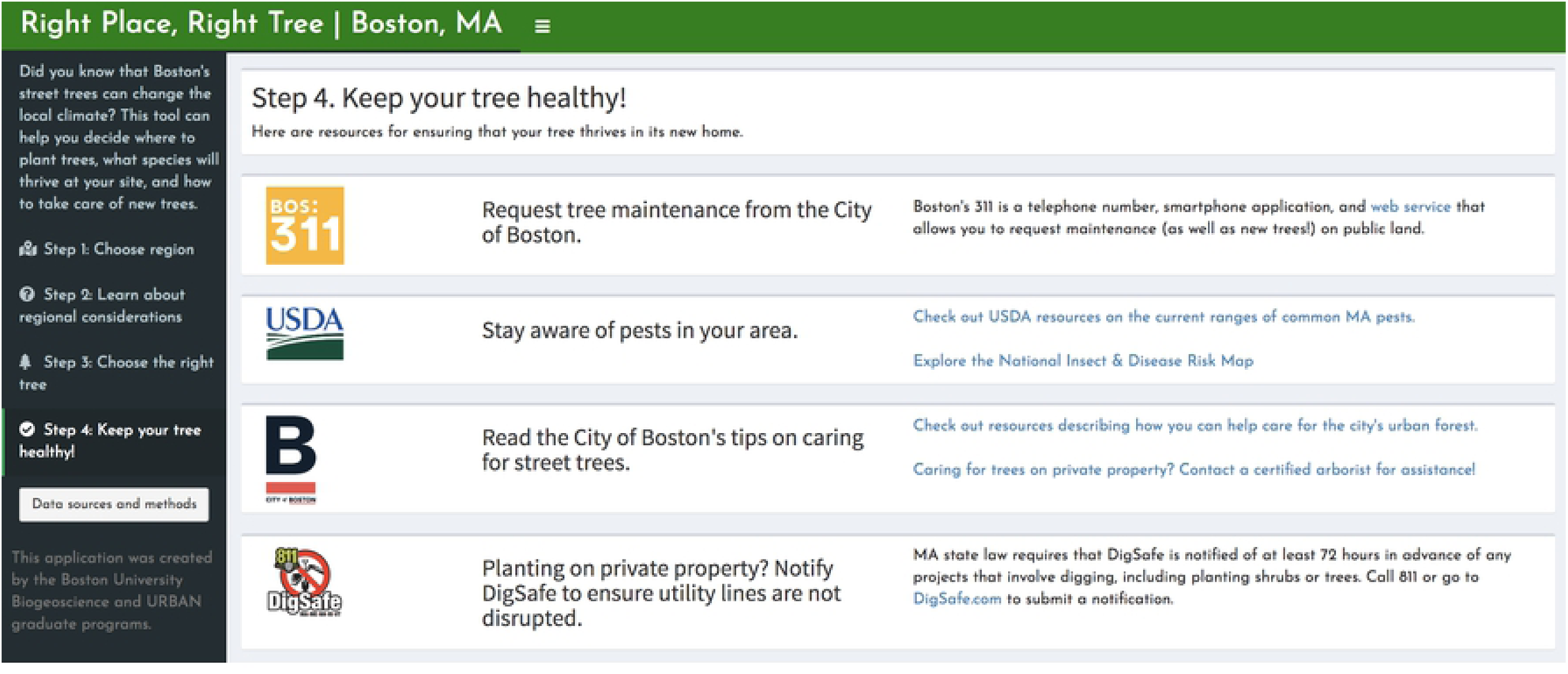
“Step 4: Keep your tree healthy.” The fourth page of the tool provides users with resources that can be consulted for maintenance of the tree they have selected using steps 1 through 3. This includes links for contacting Boston’s tree maintenance teams, up-to-date information about pests, and tips for maintenance from the Boston.gov website.

This tool was created to address the obstacles to successful canopy expansion in Boston. For instance, HVI metrics are rarely ever used for decision-making, possibly due to difficulty in interpretation [42]. Additionally, HVIs derived from larger geographies and applied to smaller areas like an individual city, have produced mixed results [14,43–45], leading to calls for HVIs tailored for specific regions [15]. One study applied a national HVI to Boston, but concluded that a Boston-specific HVI and geospatial analysis of surface temperatures was necessary [46]. We used similar methods to construct our HVI, but used data and methods to weight the factors specifically to Boston’s geography and demographic makeup. Compared to the previous HVI [46], we also included additional variables in our HVI, capturing linguistic isolation, disability and employment status, highly developed land, and age and density of housing. Another obstacle to canopy expansion has been tree maintenance and upkeep. We therefore highlight potential community partners that may be contacted for maintenance help, and the species selector can show only trees that are resistant to breakage, which may have lower maintenance requirements.

### Stakeholder Engagement

Development of this decision-making tool began in response to a request from a City of Boston representative, who outlined general questions of interest relating to Boston’s tree canopy. We found that many questions had been thoroughly answered in the scientific literature, but had not reached Boston’s tree planting efforts or prioritization. A Boston-specific HVI analysis, coupled with a new species selector tool, could help translate scientific knowledge into practice. After an initial presentation of the tool to city officials, we received feedback that the tool could assist more with the practical aspects of decision-making, so we redesigned the app to include HVI and feasibility analysis results, other related information, and a clear step-by-step decision making framework. We found that engaging meaningfully and repeatedly with the City helped ensure that our work would have a focused, direct effect on the practical decisions regarding mitigation of the urban heat island effect with tree canopy cover enhancement.

Previous research shows that canopy expansion programs can fail due to poor perception by residents, caused by (among other things) exclusion of the city residents from decision-making about tree species and maintenance [34]. The decision-making tool was therefore designed for use by city residents as well as city officials. We include information on how residents, once they have selected a tree species, can request a City-funded tree planting on public land. We also include information on obtaining and caring for healthy trees if residents plant on their own property.

### Broader Applications

Boston is just one of a handful of cities that have struggled to expand urban tree canopy cover (see discussions of Detroit and San Francisco, respectively, in [34] and [47]). We therefore designed our decision-making framework and Shiny application to be extendable for almost any U.S. city. The Supplementary Materials include methods for calculation of a HVI using census tract data in SAS. The code for the Shiny application is hosted in Github, with commentary for adapting the text and data to other cities. The species-selector tool requires aggregating tree characteristic data for regional species, which we did manually due to our variables of interest, but for a limited number of variables the i-Tree Species Utility [24] makes this task straightforward: users can download a CSV file containing data on all trees for a region. While the regional species, demographic makeup, and urban layout will differ among U.S. cities, our proposed decision-making framework and tool provides a systematic way to balance many variables.

## Conclusion

The expansion of urban canopy can provide many environmental and social benefits to a city, but planting and maintenance of trees requires thoughtful decision-making and proactive maintenance. Because the City of Boston has faced obstacles to creating lasting expansion of urban canopy, researchers from Boston University reviewed the City’s approach to canopy expansion. We identified obstacles in incorporating scientific knowledge into tree-planting decisions, and worked with the city to design a tool to facilitate decision-making. The tool itself can be replicated or adapted for other cities, and the development process serves as an example of a successful academic-public partnership. We anticipate strengthening this partnership as an active part of building climate resilience in Boston.

## Acknowledgements

We thank Dr. Alison Brizius from the City of Boston for guiding our work. We also thank the BU Biogeoscience and Environmental Health programs for facilitating meeting times and spaces. We thank Dr. Patricia Fabian and Dr. Pat Kinney for guiding the HVI work and for assistance from the Biostatistics and Epidemiology Data Analytics Center (BEDAC).

## Supporting Information

**S1 Methods. Heat Vulnerability Index data sources and calculations**.

**S1 Table. Data sources and variables used in Heat Vulnerability Index calculation**.

**S2 Table. Principal component factor weights used to calculate Heat Vulnerability Index**.

**S3 Table. Heat Vulnerability Index, summer land surface temperature, and potential tree canopy for all Boston census tracts**.

